# Ancestral duplicated *DL/CRC* orthologs display function on orchid reproductive organ innovation

**DOI:** 10.1101/2020.02.12.945865

**Authors:** You-Yi Chen, Yu-Yun Hsiao, Chung-I Li, Chuan-Ming Yeh, Nobutaka Mitsuda, Hong-xing Yang, Chi-Chou Chiu, Song-Bin Chang, Zhong-Jian Liu, Wen-Chieh Tsai

**Affiliations:** Department of Life Sciences, National Cheng Kung University, Tainan, 701, Taiwan; Institute of Tropical Plant Sciences and Microbiology, National Cheng Kung University, Tainan, 701, Taiwan; Orchid Research and Development Center, National Cheng Kung University, Tainan 701, Taiwan; Department of Statistics, National Cheng Kung University, Tainan, 701, Taiwan; Bioproduction Research Institute, National Institute of Advanced Industrial Science and Technology (AIST), Tsukuba, Ibaraki 305-8566, Japan; Division of Strategic Research and Development, Graduate School of Science and Engineering, Saitama University, Saitama, Saitama 338-8570, Japan; Shanghai Key Laboratory of Plant Functional Genomics and Resources, Chenshan Plant Science Research Center, CAS, Shanghai Chenshan Botanical Garden, Shanghai, China; Key Laboratory of National Forestry and Grassland Administration for Orchid Conservation and Utilization at College of Landscape Architecture, Fujian Agriculture and Forestry University, Fuzhou 350002, China

**Author notes:** Correspondence and requests for materials should be addressed to W.-C.T.

**Keywords:** orchid, *Phalaenopsis equestris*, YABBY gene, *DROOPING LEAF/ CRABS CLAW*, gynostemium, ovule

## Abstract

The orchid flower is renowned for complexity of flower organ morphogenesis and has attracted great interest from scientists. The YABBY genes encode plant-specific transcription factors with important roles in vegetative and reproductive development in seed plants. *DROOPING LEAF*/*CRABS CLAW* (*DL*/*CRC*) orthologs are involved in reproductive organ development (especially carpels) of angiosperms. Orchid gynostemium (the fused organ of the androecium and gynoecium) and ovule development are unique developmental processes. Understanding the *DL/CRC-like* genes controlling the developmental program of the gynostemium and ovule could provide accessible information for reproductive organ molecular regulation in orchids. Two *DL/CRC-like* genes, named *PeDL1* and *PeDL2*, were cloned from *Phalaenopsis equestris*. The orchid DL/CRC forms a monophyletic clade with two subclades including AshDL, PeDL1 and DcaDL1 in subclade I, and PeDL2 and DcaDL2 in subclade II. The temporal and spatial expression analysis indicated *PeDL* genes are specifically expressed in the gynostemium and at the early stages of ovule development. Both *PeDLs* could partially complement an *Arabidopsis crc-1* mutant. Transient overexpression of *PeDL1* in *Phalaenopsis* orchids caused abnormal development of ovule and stigmatic cavity of gynostemium. PeDL1, instead of PeDL2, could form a heterodimer with PeCIN8. Paralogue retention and subsequent divergence of the gene sequence of *PeDL1* and *PeDL2* in *P. equestris* might result in the differentiation of function and protein behaviors. These results reveal the important roles of *PeDLs* involved in orchid gynostemium and ovule development and provide new insights for further understanding the molecular mechanisms underlying orchid reproductive organ development.

## INTRODUCTION

Orchidaceae represents one of the largest families of angiosperms and is known for its complexity of flowers. Orchid flowers are composed of five whorls of three segments each, including the outermost whorl of sepals, the second whorl of two petals and a highly elaborate labellum with distinctive shapes and color patterns, the third and fourth whorls of six staminal organs, and the central whorl of fused male and female reproductive organs called the gynostemium. In addition, the abaxial side of the gynostemium forms the stigmatic cavity for deposition of pollinia. The innovated floral organs of the labellum and gynostemium make the orchid flower zygomorphic and are commonly invoked as a crucial pairing for attracting and interacting with pollinators (Rudall & Bateman, 2002). The fascinating and complex structures of both floral organs have attracted great interest from evolutionary and developmental biologists. In addition to unique floral morphology, orchids are unusual among flowering plants in that in many species the ovule is not mature at the time of pollination, and the ovule initiation is precisely triggered by the deposition of pollinia into the stigmatic cavity (Tsai *et al.*, 2008; Chen *et al.*, 2012; Dirks-Mulder *et al.*, 2019).

Over the last couple of decades, genes involved in angiosperm floral organ specification were identified and the ‘ABCDE model’ was proposed, with the combined effect of the A-, B-, C-, D-, and E-class MADS-box genes determining the floral organ identity. In monocot orchids, the *Phalaenopsis* species was used as a model plant to delineate the B-, C-, D-, and E-class functions of MADS-box genes participating in specialized floral organ development (Pan *et al.*, 2014; Tsai *et al.*, 2014). The most notable feature is that *P. equestris* contains four B-class *AP3*-like genes with differential expression largely responsible for differentiation of the two closely-spaced tripartite perianth whorls into three sepals of the outer whorl, versus the inner whorl of two lateral petals and a median labellum (Tsai *et al.*, 2004). Later, the refined ‘orchid code’ and ‘Homeotic Orchid Tepal (HOT)’ models were, respectively, proposed to illustrate the regulation of perianth morphogenesis in orchids (Mondragon-Palomino & Theissen, 2011; Pan *et al.*, 2011). The studies of floral terata from *Phalaenopsis* and *Cymbidium* agreed that C- and D-class MADS-box genes correlated with the gynostemium and ovule development, respectively (Wang *et al.*, 2011; Chen *et al.*, 2012). In addition, a comprehensive study on the E-class MADS-box genes in *P. equestris* revealed that PeSEP proteins have important competence for complex maintenance, and the outcomes of the interactions are essential to regulate varied orchid tepal morphogenesis (Pan *et al.*, 2014). Recently, it has been reported that *AGL6*-like MADS-box genes have functions related to labellum development, and differential interactions among *AGL6*-like genes and *AP3*-like genes are required for sepal, petal, and labellum formation (Hsiao *et al.*, 2013; Hsu *et al.*, 2015).

The YABBY (YAB) gene family specifies the adaxial-abaxial leaf polarity (Bowman, 2000). In *Arabidopsis*, the plant-specific transcriptional factor YAB family contains six subfamilies: the FILAMENTOUS FLOWER (FIL) subfamily, the YAB2 subfamily, the YAB3 subfamily, the YAB5 subfamily, the DROOPING LEAF/CRABS CLAW (DL/CRC) subfamily and the INNER NO OUTER (INO) subfamily, which encode proteins possessing a typical zinc finger domain and a YAB domain (Bowman & Smyth, 1999; Sawa *et al.*, 1999; Siegfried *et al.*, 1999; Villanueva *et al.*, 1999). Previous studies indicated that the *Arabidopsis CRC* is specifically expressed at the carpel and plays an important role in carpel development and determinacy of the floral meristem (Alvarez & Smyth, 1999; Bowman & Smyth, 1999). The rice *dl* mutants showed that drooping leaf phenotypes resulted from lacking a leaf midrib. In addition, the mutants also demonstrated that carpels homeotically transformed into stamens, indicating that *DL* regulates carpel identity (Nagasawa *et al.*, 2003). It also indicated that *DL* interacts antagonistically with class B genes and controls floral meristem determinacy (Yamaguchi *et al.*, 2004). Ancestral expression patterns and functional studies performed with *Petunia hybrida*, *Nicotiana tabacum* and *Pisum sativum* support an ancestral role of *DL/CRC* genes in the specification of carpel development and floral meristem termination (Lee *et al.*, 2005; Yamada *et al.*, 2011; Fourquin *et al.*, 2014). Interestingly, in basal eudicot California poppy, *EcCRC* not only harbors ancestral functions of *DL/CRC* but is also involved in ovule initiation (Orashakova *et al.*, 2009).

In this study, we identified eight *YAB* genes, including two members in the CRC clade, one member in the INO clade, three members in the YAB2 clade and two members in the FIL clade, from the *P. equestris* whole genome sequence (Cai *et al.*, 2015). We determined *PeDL*s expression patterns and performed transgenic overexpression of *PeDL*s in wild-type *Arabidopsis* and a *crc* mutant, and transient overexpression in *Phalaenopsis* orchids to functional characterization of *PeDL*s. Potential candidate targets of *PeDL*s were revealed by transcriptomic comparison. We concluded that *PeDL*s not only have conserved function on floral meristem determinacy and carpel development, but also have novel function on stigmatic cavity formation and ovule initiation.

## RESULTS

### Identification and phylogenetic analysis of orchid YABBY genes

The *P. equestris*, *D. catenatum*, and *A. shenzhenica* genomes respectively encode 8, 7, and 5 YAB genes. The number of YAB genes of *P. equestris* and *D. catenatum* is comparable to that of *Arabidopsis* (6 members) and rice (8 members). The multiple sequence alignment analysis showed that protein sequences translated by *P. equestris’* eight YAB genes harbor standard zinc finger and YAB domains (Fig. 1b, c) (Bowman & Smyth, 1999). To determine phylogenetic relationships among the orchid and other plant YAB genes, the phylogeny of known YAB genes was reconstructed using the conceptual amino acid sequences of respective genes as input data. The topology of the phylogenetic tree obtained indicated that the orchid and rice genes reported here fell well into the four clades including the YAB2, DL/CRC, INO and FIL with use of gymnosperm YAB genes as an outgroup (Fig. 1a) (Finet *et al.*, 2016). The results suggested that the last common ancestor of monocots might loss the YAB5 subfamily (Fig. 1a). Furthermore, PeYAB1 and PeFIL belong to the FIL clade; PeYAB2, PeYAB3 and PeYAB4 are classified into the YAB2 clade; PeINO is in the INO clade; and PeDL1 and PeDL2 are members of the DL/CRC clade (Fig. 1a).

**Fig. 1.**
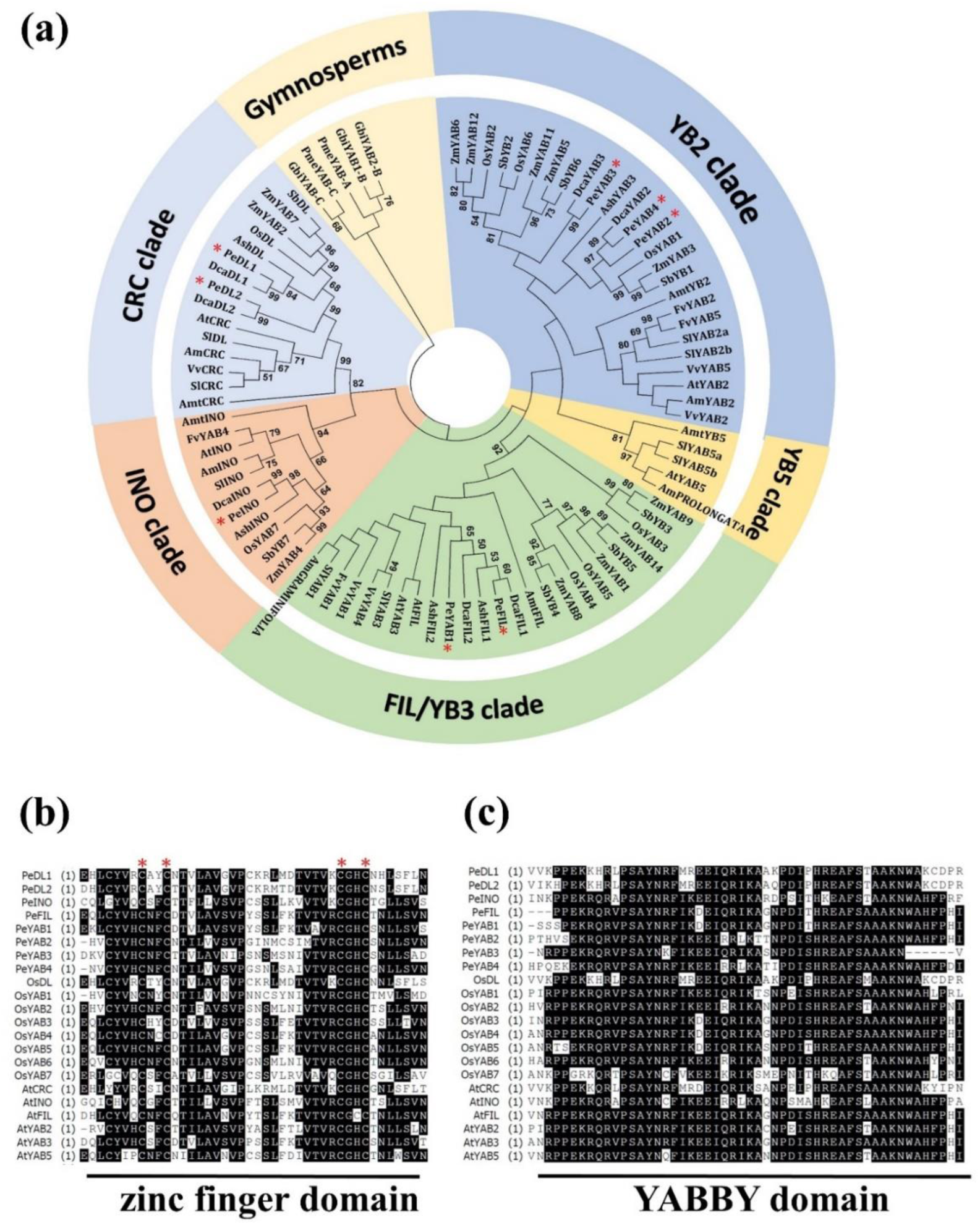
Sequence analysis of plant YABBY genes. (a) Phylogenetic analysis of YABBY-related proteins from the angiosperms and gymnosperms. The PeYABBY-related proteins were marked by red asterisks. The tree was generated using MEGA6.0 neighbor-joining software with 1000 bootstrap trials. Accession numbers are listed in Table S1. (b) Protein sequence alignment of the C2C2 zinc finger domain and YABBY domain from rice, *Arabidopsis* and *P. equestris*. Consensus is denoted by the color black; similarity is denoted by gray. Conserved cysteine residues in the zinc-finger domain are indicated with red asterisks.

It has been indicated that the DL/CRC subfamily represents a single orthologous lineage, without ancient duplications (Lee *et al.*, 2005). Interestingly, we found that both *P. equestris* and *D. catenatum* have two *DL/CRC*-like genes and only one was identified in *Apostasia* (Fig. 1). We further compared the gene structures of *DL/CRC* genes among *P. equestris*, *A. shenzhenica*, *D. catenatum*, *Cabomba caroliniana*, *Oryza sativa* and *A. thaliana*. All the *DL/CRC* genes have one intron located in the zinc finger domain, three introns located in the YAB domain and one intron situated in the non-conserved region between the zinc finger domain and YAB domain (Fig. S1a and S1b). Interestingly, the sixth intron could be found in the region downstream of the YAB domain in most of the plants, except *C. caroliniana CcCRC* and *D. catenatum DcaDL2*. The multiple alignments of protein sequences showed that both PeDL1 and PeDL2 contain the typical zinc finger domain and YAB domain which are consistent with other plant species with DL/CRC orthologous proteins (Fig. S1c). The phylogenetic tree showed that orchid DL/CRC form a monophyletic clade with two subclades, including AshDL, PeDL1 and DcaDL1 in subclade I, and PeDL2 and DcaDL2 in subclade II (Fig. S1d). Paralogue retention and subsequent divergence of gene sequences of the *DL/CRC*-like genes in *P. equestris* and *D. catenatum* might result in the functional differentiation of the *DL/CRC*-like genes.

### Expression profiles of *PeDL* genes in *P. aphrodite* subsp. *formosana*

The presence of two *DL*/*CRC*-like paralogs in *Phalaenopsis* raises the question whether the two genes are functionally redundant or are regulated temporally and spatially during reproductive organ development. To distinguish between these two possibilities, spatial and temporal expressions of the two *PeDL* paralogs were investigated by quantitative real-time RT-PCR (Fig. 3a-f). The results showed that both *PeDL1* and *PeDL2* were predominantly expressed at early flower bud developmental stages (Fig. 2a, 2b, 3a, and 3b). In addition, transcripts of both *PeDL1* and *PeDL2* could be specifically detected in gynostemium (Fig. 2c, 3c, and 3d). The expression of these two genes was not detectable in vegetative tissues, such as pedicels, floral stalks, leaves and roots (Fig. 2b, 2d, 2e, 3c, and 3d). According to the highly consistent spatial and temporal expression patterns in flowers of these two *PeDL* genes, it is possible that both *PeDL* genes have important functions for gynostemium development.

**Fig. 2.**
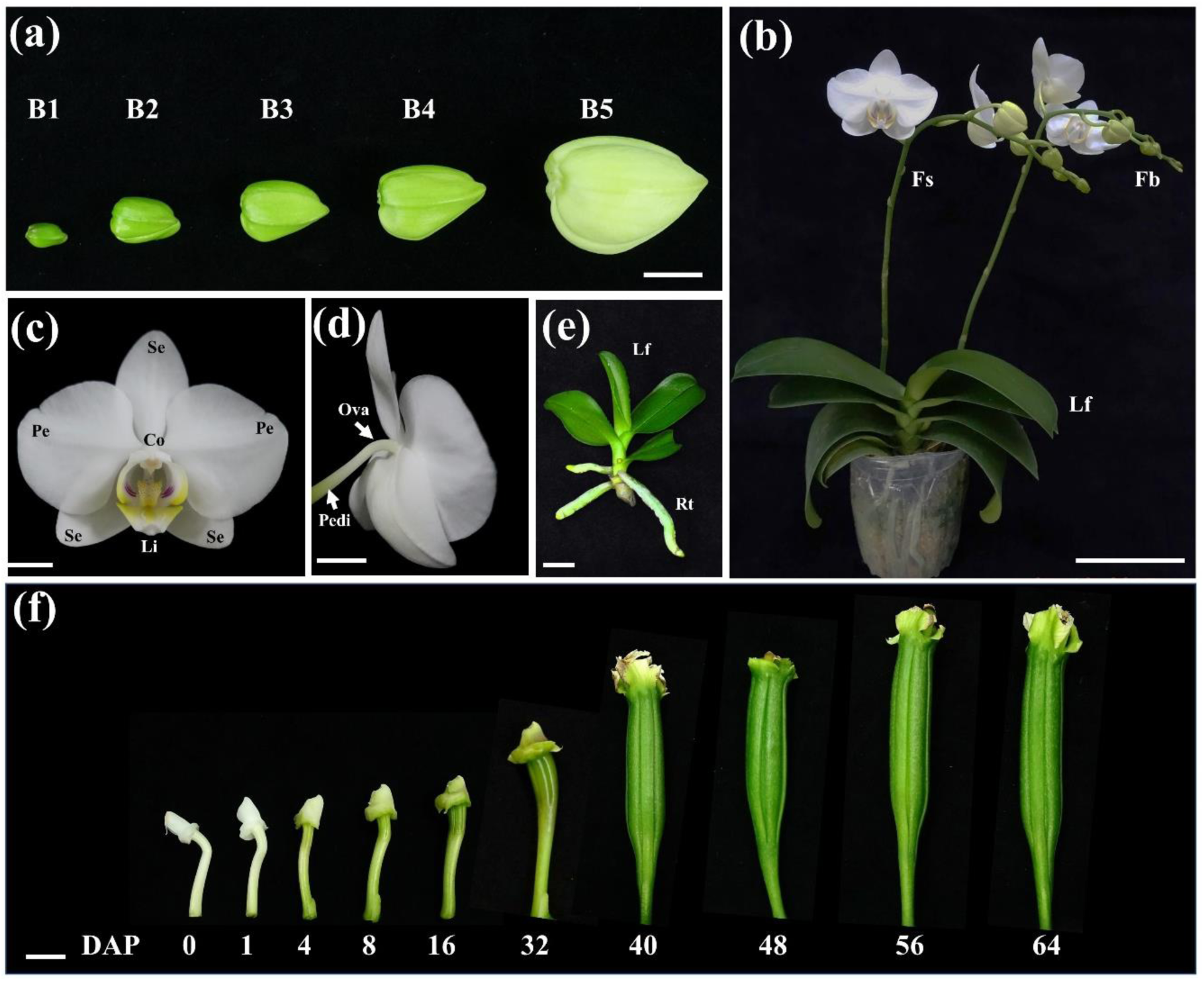
*P. aphrodite* subsp. *formosana*. (a) Flower bud development stages from B1 to B5, Scale bar = 1 cm; (b) Flower buds, fully blooming flowers and floral stalk in mature *Phalaenopsis* plant. Scale bar = 10 cm; (c, d) Structure of *P. aphrodite* subsp. *formosana* flower. Scale bar =1cm; (e) Leaf and root at the vegetative seeding stage. Scale bar = 1 cm; (f) Morphological changes in the ovary at various days after pollination. Scale bar = 1cm. Se, sepals; Pe, petals; Li, lip; Co, column (gynostemium); Pedi, pedicel; Fs, floral stalk; Lf, leaf; Rt, root; Ova, ovary; DAP, days after pollination.

**Fig. 3.**
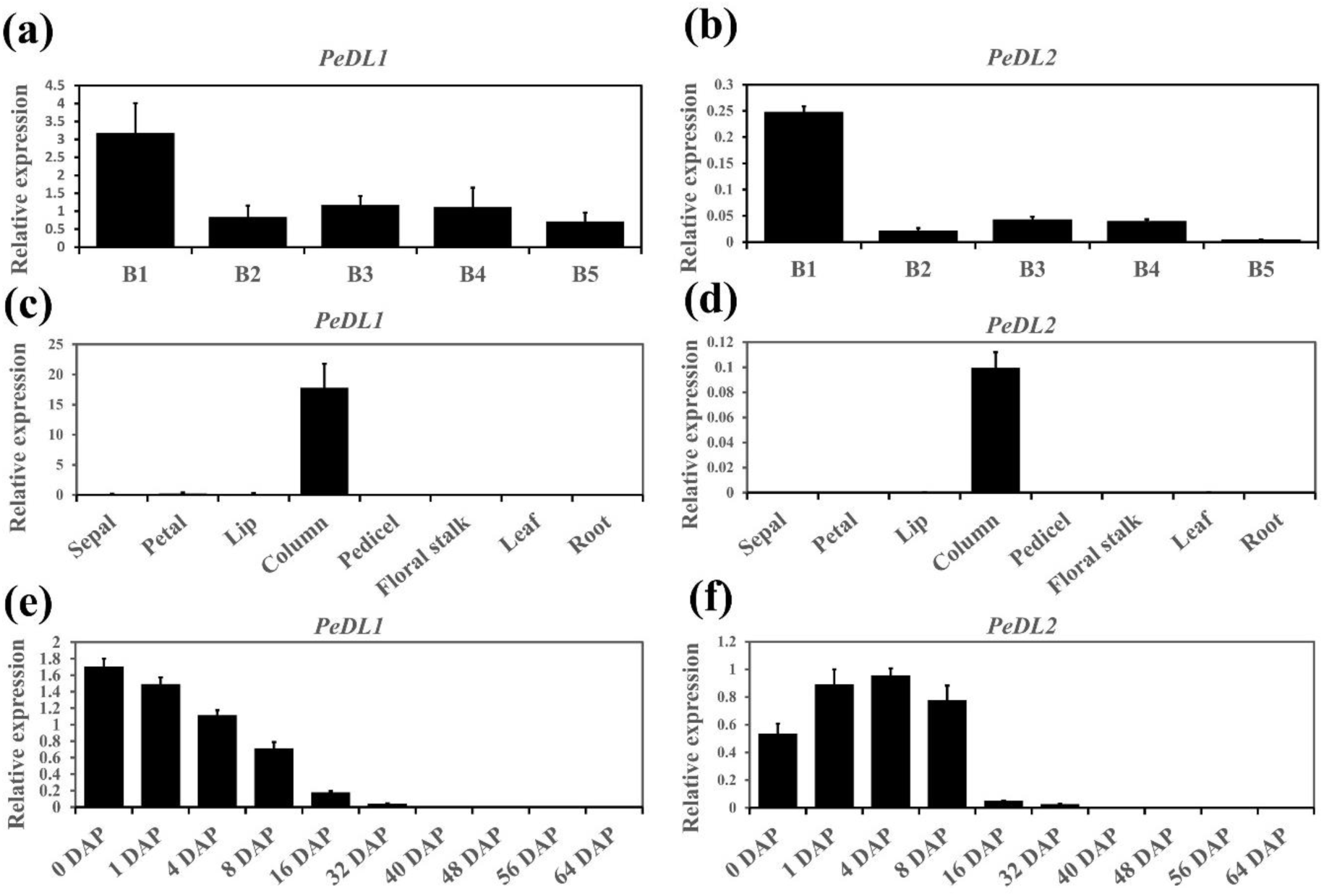
Analyses of spatial and temporal expression patterns of *PeDL1* and *PeDL2* in *P. aphrodite* subsp. *formosana* by quantitative real-time PCR. (a, b) Expression patterns of *PeDL1* and *PeDL2* in developmental stages of flower buds. B1, stage 1 flower buds (0.5–1 cm); B2, stage 2 flower buds (1–1.5 cm); B3, stage 3 flower buds (1.5–2 cm); B4, stage 4 flower buds (2–2.5 cm); B5, stage 5 flower buds (2.5–3 cm). (c, d) Expression patterns of *PeDL1* and *PeDL2* in various plant organs; (e, f) Expression patterns of *PeDL1* and *PeDL2* in various ovule developmental stages. DAP, days after pollination.

Because YAB genes are also involved in ovule development, and pollination is a key regulatory event in orchid ovule initiation, we determined the temporal mRNA expression patterns of both *PeDL*s in developing ovules triggered by pollination (Fig. 2f). Both expressions of *PeDL1* and *PeDL2* could be detected before 32 days after pollination (DAP). After pollination, the expression of *PeDL1* gradually decreased to 32 DAP (Fig. 3e). However, the expression of *PeDL2* gradually increased to 4 DAP, and then decreased, and then was not detectable after 32 DAP (Fig. 3f). Therefore, both of these *PeDL*s’ functions may also be associated with ovule development. Based on the substantially differential expressions of the two *PeDL* paralogs, it is possible that they have functional differentiation in ovule initiation.

The detailed spatial and temporal expression patterns of the *PeDL*s gene in gynostemium and ovule development was investigated by means of *in situ* hybridization by using antisense RNA as a probe. Both *PeDLs* transcripts were detected in the inflorescence meristem and floral primordia (Fig. 4a and S2a). At the early flower bud stage, transcripts of *PeDL1* and *PeDL2* could be strongly detected in the gynostemium (column) and column foot, the basal protruding part of the column (Fig. 4e and S2e). At the later developmental stages, the transcripts of *PeDL1* were more strongly concentrated at the rostellum, the projecting part of the column that separates the male androecium from the female gynoecium, and the column foot (Fig. 4c and S2c). In the process of ovule development, *PeDL* transcripts were detected in the pericarp and placenta before pollination (Fig. 4g and S2g). At 4 and 8 DAP, both *PeDL*s were expressed in the pericarp and developing ovules (Fig. 4i, 4k, S2i and S2k). At 16 DAP, the signals of both *PeDL*s could be detected in developing ovules and the first layer cells of the placenta (Fig. 4m and S2m). The results of expression analysis suggested that both *PeDL* genes might be involved in gynostemium development as well as ovule initiation.

**Fig. 4.**
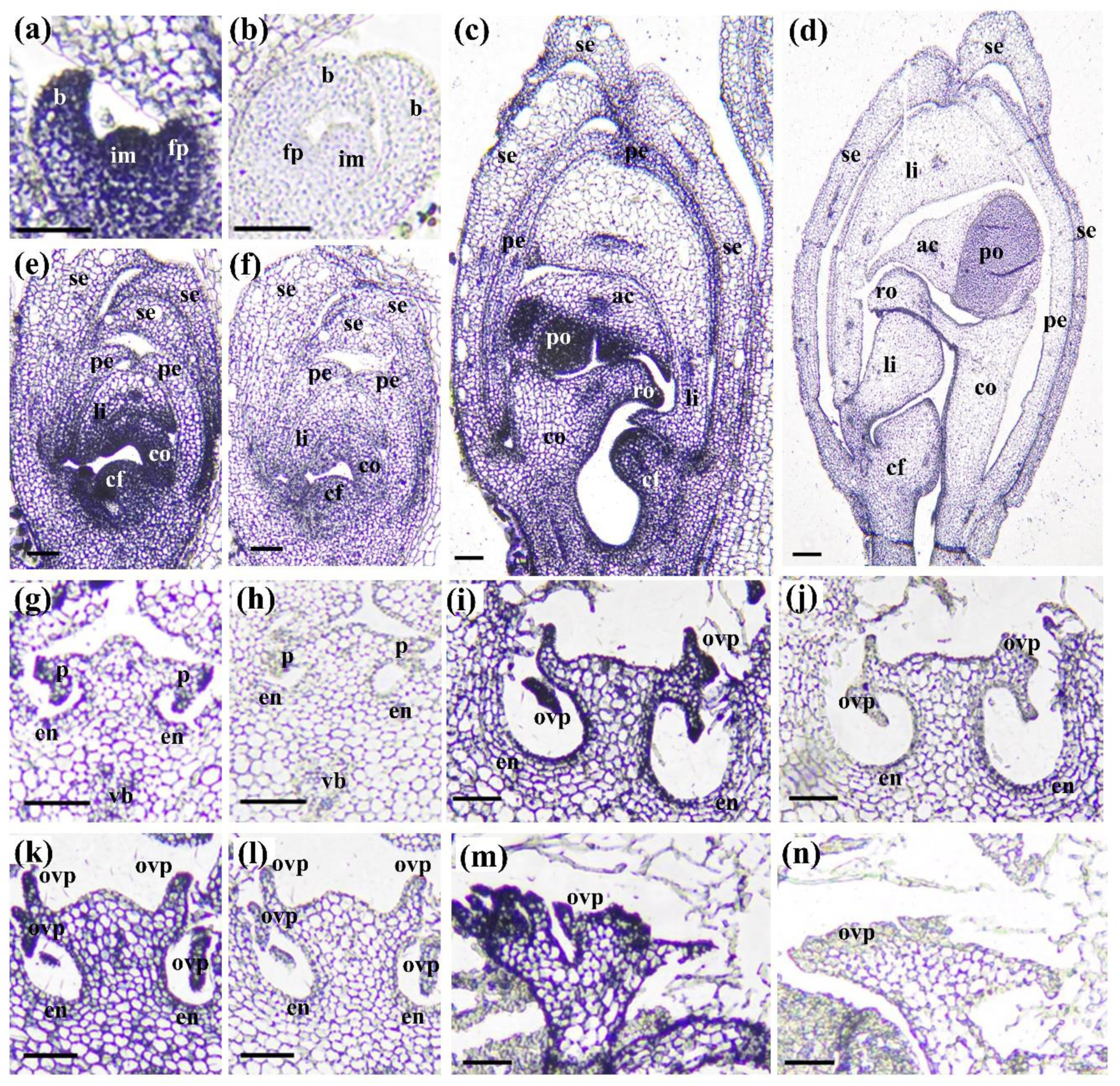
RNA *in situ* hybridization of *PeDL1* longitudinal sections in developing floral buds and cross sections in developing ovules of *P. equestris*. In panels (a, c, e, g, i, k, m), antisense probes were used to detect *PeDL1* transcripts. In panels (b, d, f, h, j, l, n), hybridization was done with *PeDL1* sense probes (negative controls). Sections were hybridized with the antisense 3’-specific *PeDL1* RNA probes or sense RNA probes. (a, b) Inflorescence meristem and floral primordia; (e, f) longitudinal section of the flower buds at an early stage; (c, d) longitudinal section of the flower buds at a late stage; (g, h) ovary tissue before pollination; (k, l) placenta with ovule primordia at 4 DAP; (i, j) placenta with ovule primordia at 8 DAP; (m, n) placenta with ovule primordia at 16 DAP. im, inflorescence meristem; fp, floral primordia; se, sepal; pe, petal; li, lip; co, column; cf, column foot; ro, rostellum; ac, anther cap; po, pollinium; b, bract; ovp, ovule primordia; p, placenta; en, endocarp; vb, vascular bundles. (a-n) Bar = 100 µm.

### Functional analyses of the two *PeDL* genes using transgenic *Arabidopsis*

To investigate the role of the putative function of *PeDL* genes, we overexpressed *PeDL1* and *PeDL2* using the *Agrobacterium*-mediated method under control of the *Cauliflower mosaic* virus (CaMV) *35S* promoter in *Arabidopsis*, respectively. Both overexpressed plants showed reduced plant size, and *35S::PeDL1* produced smaller plants than *35S::PeDL2* (Fig. S3a). In addition, both transgenic plants produced rosette leaves curled towards the abaxial side with wrinkled surfaces (Fig. S3a). Epidermal cell observations revealed that in both *35::PeDL1* and *35::PeDL2* epidermal cell size was decreased (Fig. S3b, 3c, 3d, and 3h), along with increased cell density in the adaxial side of rosette leaves (Fig. S3c, 3d, and 3i). However, no significant changes in abaxial epidermal cells were observed among wild type (WT) and transgenic plants (Fig. S3e, 3f, 3g, and 3i).

Both overexpressed plants showed short siliques containing reduced numbers of seeds as compared to those of WT plants (Fig. 5a, 5e-5j). The seed size of *35S::PeDL1* plants was larger than WT plants and the seed weight was significantly increased (Fig. S4). Interestingly, both *35S::PeDL1* and *35S::PeDL2* plants presented shriveled styles and replum, and the style cell size was smaller than that of WT (Fig. 5b, 5c, and 5d). These observed gynoecium abnormalities of overexpressed plants suggested that *PeDL1* and *PeDL2* are important for gynoecium development. We also observed that the primary inflorescence apices were abnormally terminated and showed a cluster of flower buds in *35S::PeDL1* and *35S::PeDL2* plants (Fig. S5a-S5d). This phenotype implies that *PeDL1* and *PeDL2* might be involved in termination of the primary inflorescence meristem. Previous studies also indicated that *DL*/*CRC* genes are involved in floral meristem determinacy (Alvarez & Smyth, 1999; Li *et al.*, 2011).

**Fig. 5.**
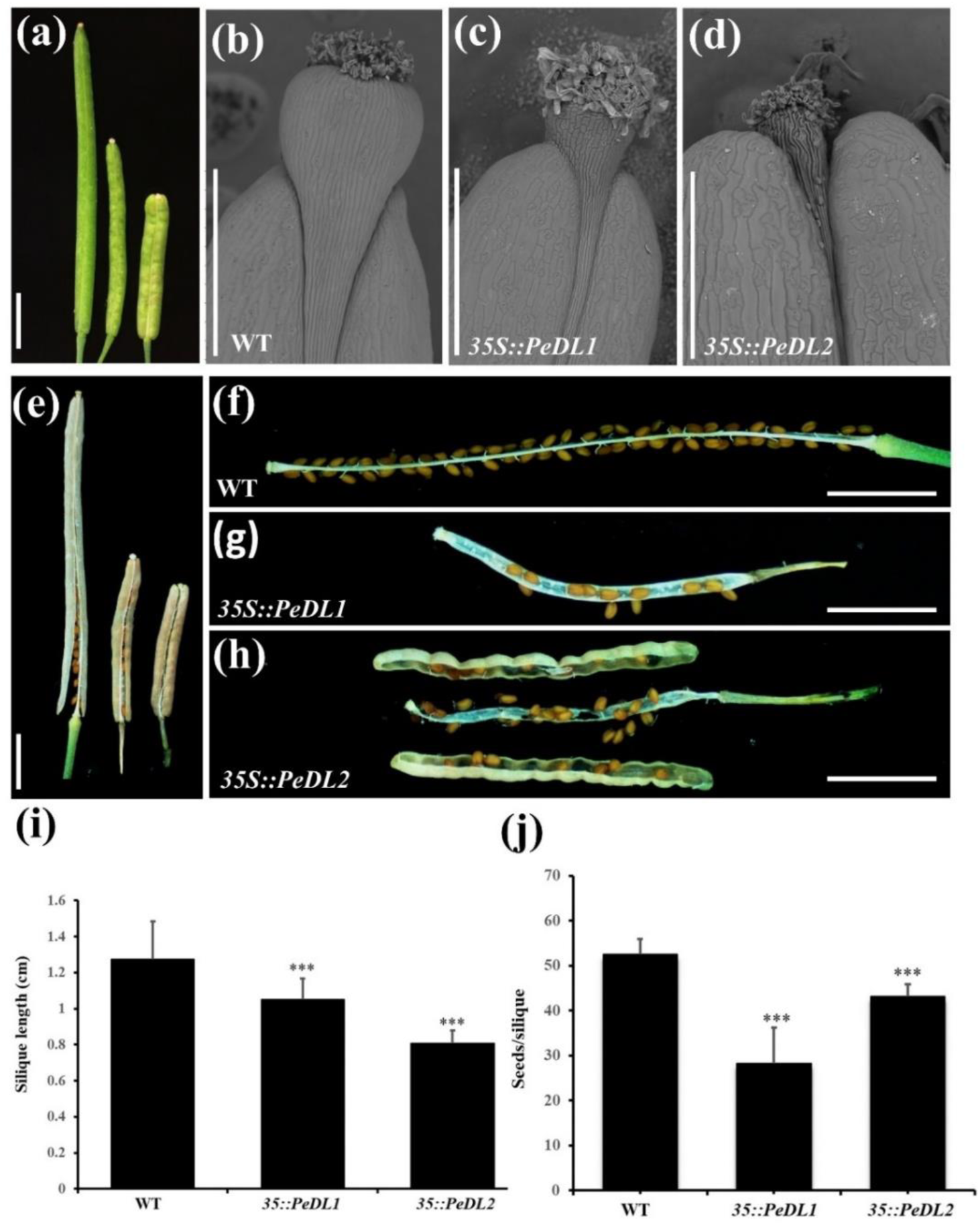
Morphology of siliques in WT, *35S::PeDL1* and *35S::PeDL2* transgenic plants. (a) Comparison of immature siliques in WT (left), *35S::PeDL1*(middle) and *35S::PeDL2*(right) transgenic plants. Scale bar= 0.25 cm; <u>S</u>canning <u>E</u>lectron <u>M</u>icrographs (SEM) of (b) WT, (c) *35S::PeDL1* and (d) *35S::PeDL2* transgenic *Arabidopsis*. Scale bar=500 µm; (e) Comparison of mature siliques in WT (left), *35S::PeDL1*(middle) and *35S::PeDL2*(right) transgenic plants. Scale bar= 0.25 cm; (f–h) Siliques of (f)WT, (g)*35S::PeDL1* and (h)*35S::PeDL2*. (f, g) with valves removed. Scale bar= 0.25 cm. (i) Mean number of silique length in WT, *35S::PeDL1* and *35S::PeDL2*. Asterisks indicate statistically significant differences (*** P<0.001, compared with WT by Student’s t-test). Error bars represent the SD of three biological repeats (n=20 each). (j) Mean number of seeds per silique in WT, *35S::PeDL1* and *35S::PeDL2*. Asterisks indicate statistically significant differences (***P<0.001, compared with WT by Student’s t-test). Error bars represent the SD of three biological repeats (n=20 each).

To further investigate *PeDL* genes’ functions, we overexpressed *PeDL1* and *PeDL2* in *crc-1* plants, respectively. The results revealed that both *PeDL* genes could partially rescue the *crc-1* mutant. The *crc-1* mutant flower is similar to that of WT plants, except the carpels are unfused at the apical region (Fig. 6a, 6b, 6c, 6f, and 6g) (Bowman & Smyth, 1999). Respective overexpression of *PeDL1* and *PeDL2* in *crc-1* plants dramatically influenced the gynoecium development. The carpels of *35S:PeDL1 crc-1* plants showed smaller gaps at the apical region of the gynoecium than that of WT plants (Fig. 6a, 6d, and 6h). The *35S:PeDL2 crc-1* plants presented perfect fused carpels although the styles and replum shriveled severely (Fig. 6a, 6e, and 6i). These results suggest that the *PeDLs* play a conserved role in controlling carpel development as eudicot *CRC*.

**Fig. 6.**
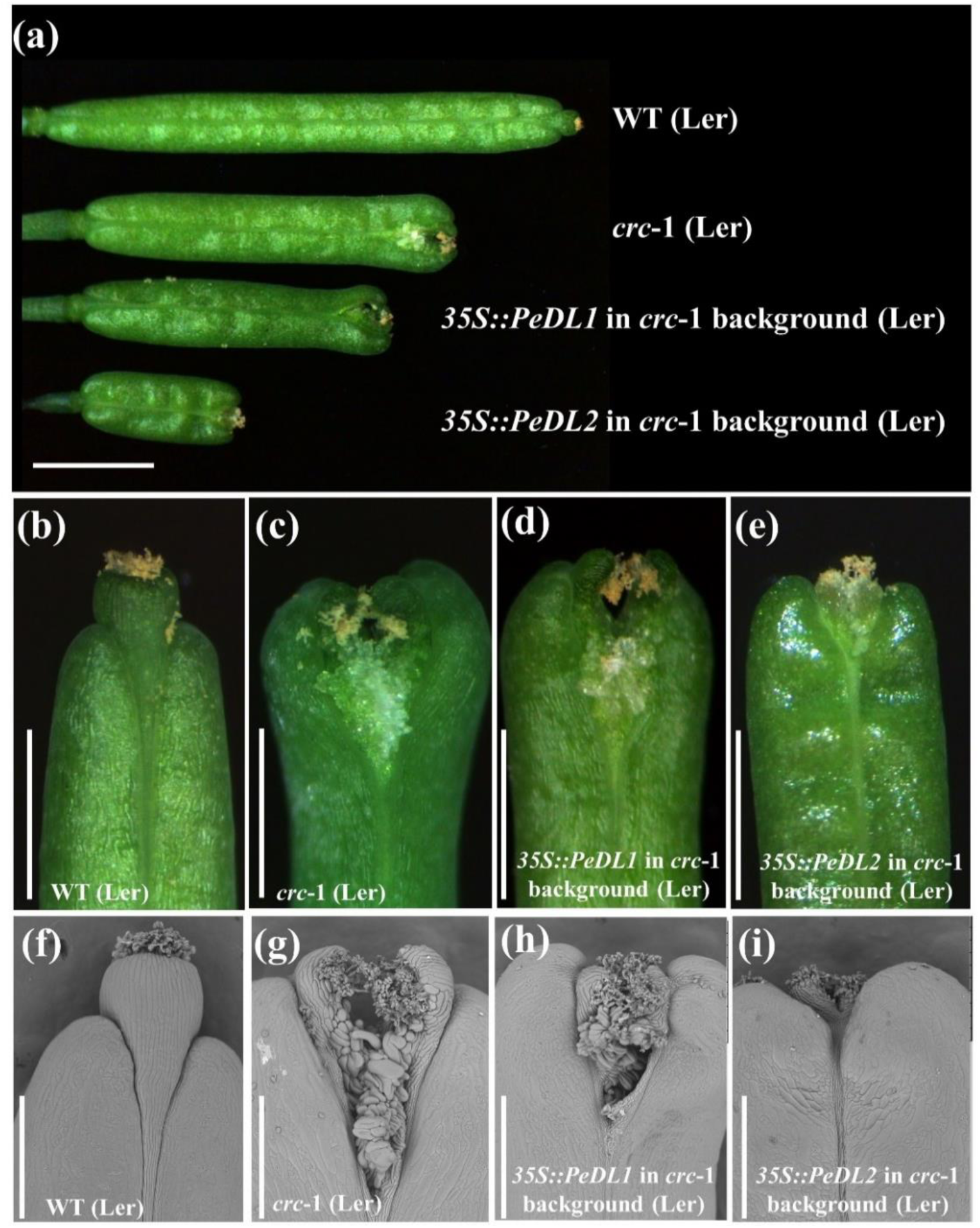
Ectopic expression of *PeDL1* and *PeDL2* in the *crc*-1 mutant respectively. (a) Comparison of immature siliques in WT, *crc-1*, overexpression *35S-PeDL1*/ *crc*-1 and overexpression *35S-PeDL2*/ *crc*-1 transgenic plants. Scale bar= 0.5 cm; (b) immature siliques of a WT plant. (c) *crc*-1 mutant immature silique. (d) immature siliques of a transgenic *crc*-1 mutant plant ectopically expressing *PeDL1*. (e) immature siliques of a transgenic *crc*-1 mutant plant ectopically expressing *PeDL2*. (f) SEM of WT immature silique. (g) SEM of *crc*-1 mutant immature silique. (h) SEM of immature siliques of a transgenic *crc*-1 mutant plant ectopically expressing *PeDL1*. (i) SEM of immature siliques of a transgenic *crc*-1 mutant plant ectopically expressing *PeDL2*. (b–e) Scale bar = 100 µm; (f–i) Scale bar = 500 µm.

### Transient overexpression of *PeDL* genes in *Phalaenopsis* orchid

To characterize the function of *PeDL*s in *Phalaenopsis* orchids, we performed transient overexpression of *PeDL* genes in *P. Sogo Yukidian*. Twenty transient overexpressions of *PeDL2* (*OE-PeDL2*) did not cause obvious phenotype changes. Seventeen plants with *PeDL1* transient overexpression (*OE-PeDL1*) were obtained. All the plants showed abnormal stigmatic cavities with variant severity. Among these, three plants with strong, six with moderate, and eight with weak severity of stigmatic cavity morphology were observed. The stigmatic cavities with strong severity showed that a protruding tissue grew from the bottom of the cavity and almost covered the cavity (Fig. 7f). The stigmatic cavities with moderate severity had a sharp (Fig. 7j) or flat tissue (Fig. 7g and S6b) extending from the bottom of the cavity. The ones with weak severity presented unapparent v-shapes (Fig. 7h, 7k, and S6c) at the bottom of the stigmatic cavity as compared to the v-shapes of mock plants (Fig. 7e, 7i, and S6a). The expression level of *PeDL1* in *OE-PeDL1* was positively correlated with the severity phenotype of the stigmatic cavity (*OE-PeDL1-2*, strong severity; *OE-PeDL1-4*, moderate severity; *OE-PeDL1-5*, weak severity), suggesting these phenotypes are dosage dependent. The flower size and the morphology of sterile floral organs were similar between mock and *OE-PeDL1* plants (Fig. 7a-7d). These results suggest that the function of *PeDL1* is associated with the morphogenesis of the stigmatic cavity.

**Fig. 7.**
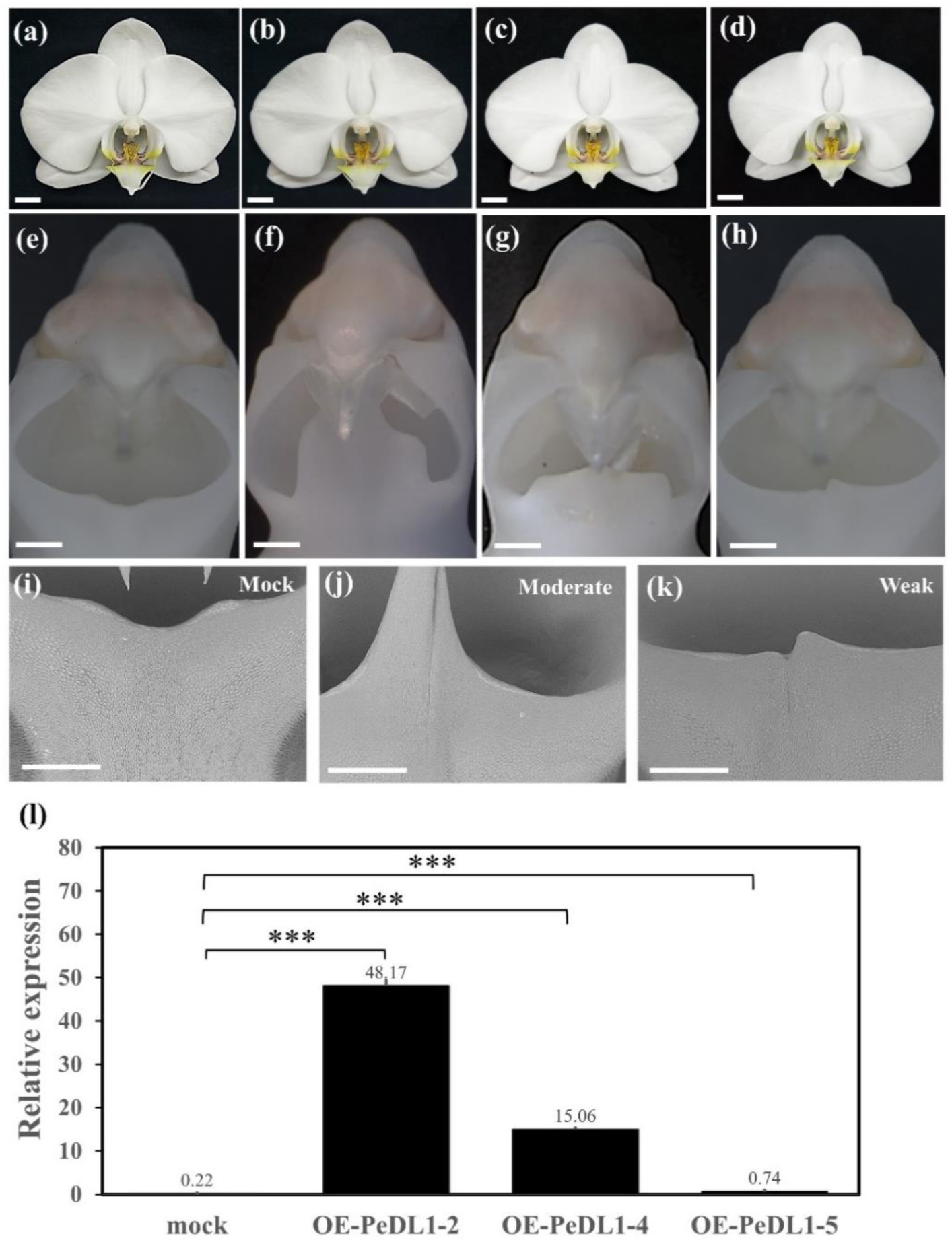
Transient overexpression of *PeDL1* in *P.* Sogo Yukidian. (a) The mock-treated flower. (b) The OE-*PeDL1*-2 flower. (c) The OE-*PeDL1*-4 flower. (d) The OE-*PeDL1*-5 flower. (e) The mock-treated gynostemium. (f) The OE-*PeDL1*-2 gynostemium (strong). (g) The OE-*PeDL1*-4 gynostemium (moderate). (h) The OE-*PeDL1*-5 gynostemium (weak). SEM of the moderate and weak phenotypes on the gynostemium of (i) mock-treated and (j, k) OE-*PeDL1* plants. (l) Relative expression of *PeDL1* mRNA level in mock-treated and *OE-PeDL1* plants. (a–d) Scale bar = 2 cm, (e–h) Scale bar = 0.1 cm, (i–k) Scale bar = 1 mm. (l) Asterisks indicate statistically significant differences (***P<0.001, compared with mock-treated by Student’s t-test).

### Transcriptome comparison between *OE-PeDL1* and mock plants

To identify the gynostemium-expressed genes influenced by the expression of *PeDL1*, we compared the gynostemium transcriptomes between *OE-PeDL1* and mock plants. The genes with expression fold-change greater than 2 and an adjusted *P*-value less than 0.05 were selected as the differentially expressed genes (DEGs). Seventy-five up-regulated (Table S2) and 180 down-regulated genes (Table S3) were identified in overexpressed *PeDL1* plants (Fig. S8). The green dot with a red circle in the volcano plot indicated that *PeDL1* had the highest expression level with significant differential expression among the 75 up-regulated genes (Fig. S8). Among 75 up-regulated DEGs, 3 DEGs including PAXXG113920 (cyclin-dependent kinase B2-1-like), PAXXG079290 (cyclin), and PAXXG072210 (cyclin-A1-4) are related to cell proliferation (Table S2). Up-regulated cell cycle related genes discovered in *OE-PeDL1* plants might explain the generation of protruding tissue growing from the bottom of the stigmatic cavity in *OE-PeDL1* plants. The results also suggest that *PeDL1* could directly or indirectly be involved in cell cycle regulation. Interestingly, five up-regulated genes, PAXXG134630, PAXXG177570, PAXXG177580, PAXXG177760, and PAXXG339360, belonging to the SUGARS WILL EVENTUALLY BE EXPORTED TRANSPORTER (SWEET) family were also identified, which might be associated with the sticky surface of the stigmatic cavity.

### *PeDL1* influences endocarp development and ovule initiation

One of the remarkable reproductive characteristics of most orchid species is that ovary and ovule development are precisely initiated following pollination (Tsai *et al.*, 2008). Interestingly, *OE-PeDL1* plants showed that ovary growth was affected. Before pollination, the ovary morphology was not significantly different between *OE-PeDL1* and mock plants (Fig. S7a-S7d). However, SEM observations showed that several empty spaces existed among three lobes of the endocarp in *OE-PeDL1* (Fig. 8b and 8d). In contrast, the mock plants presented almost completely filled endocarps (Fig. 9a and 9c). At 16 DAP, the ovaries of *OE*-*PeDL1* did not obviously enlarge as compared to those of mock plants (Fig. S7e and Fig. S7f). The number of protuberant ovule initials differentiated from the placenta was obviously decreased in *OE-PeDL1* (Fig. 8e-8h). These results suggest that *PeDL1* is involved in endocarp development and ovule initiation in *Phalaenopsis* orchids.

**Fig. 8.**
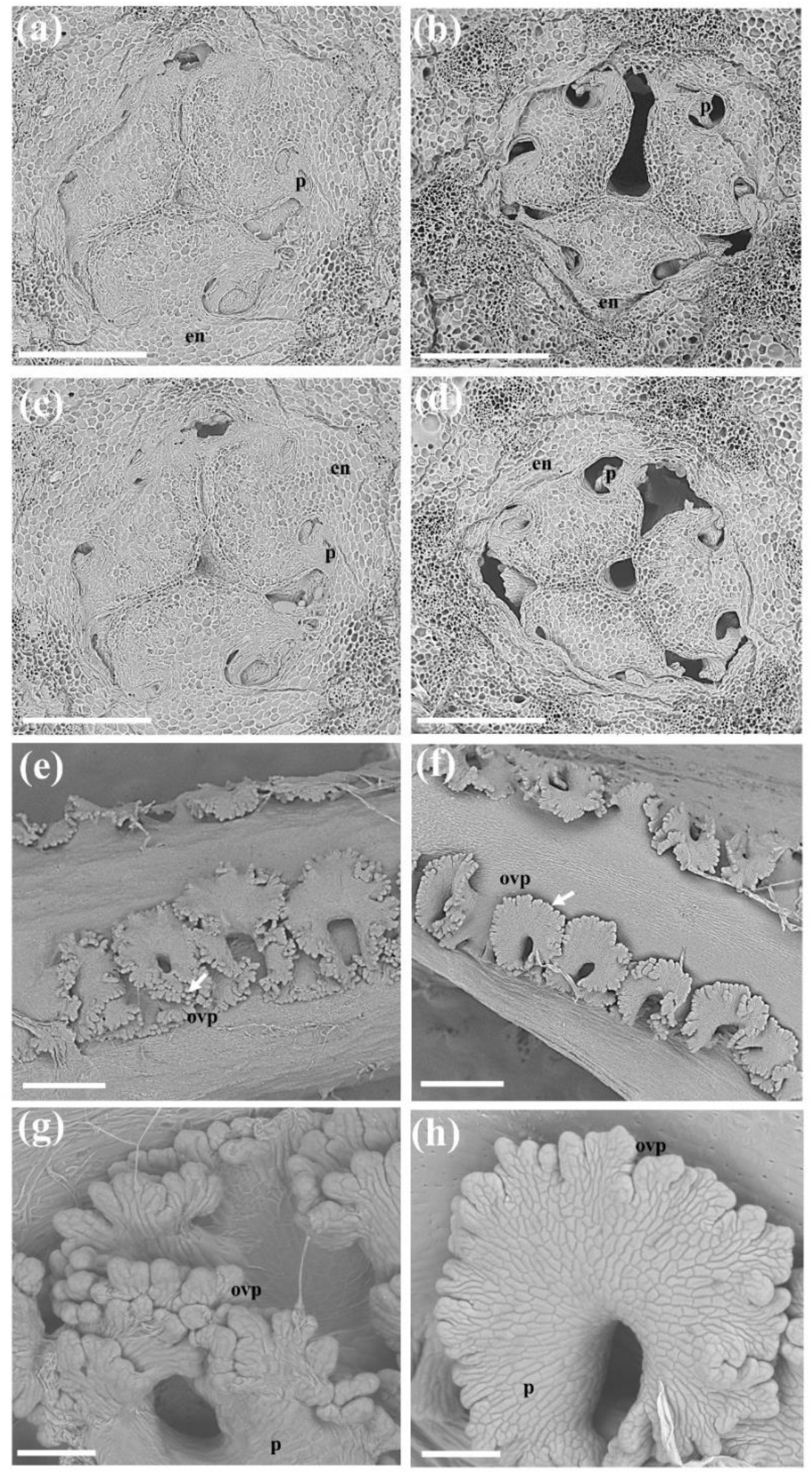
Phenotypic effects of transient overexpression *PeDL1* on ovule and placenta development of the ovary before pollination and 16 days after pollination. SEM of the (a and c) mock-treated and (b and d) OE-*PeDL1* cross section of the ovary before pollination. The (b and d) shows the shriveled placenta. SEM of (e) mock-treated and (f) OE-*PeDL1* of the 16 days after pollination. (g) Enlarged region of the white arrow in panel (e). (h) Enlarged region of the white arrow in panel (f). The (f and h) reduced development of ovule primordia. ovp, ovule primordia; p, placenta; en, endocarp.

**Fig. 9.**
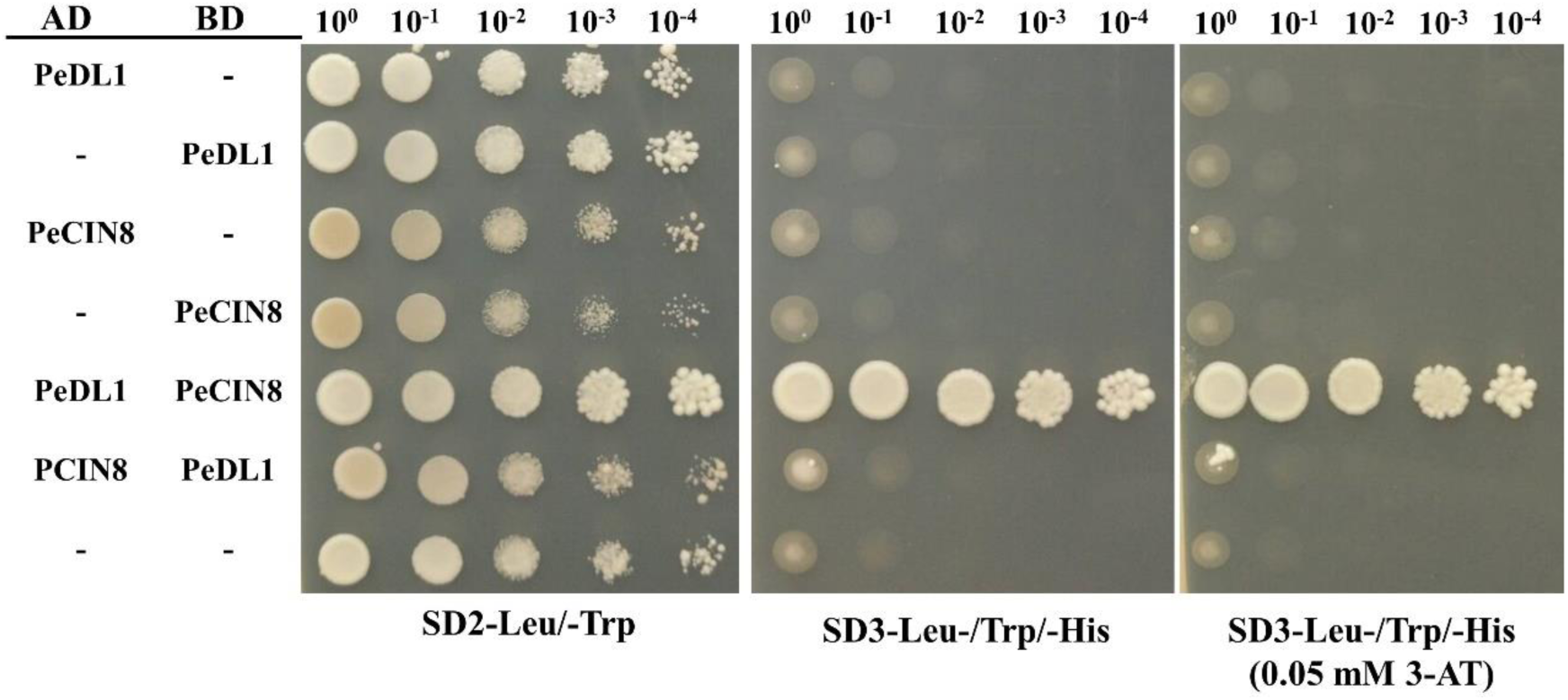
PeDL1 interacts with PeCIN8 in the yeast two-hybrid assay. PeDL1 and the PeCIN8 gene were cloned into the binding domain vector pGBKT7(BD) and the activation domain vector pGADT7(AD). Yeast AH109 strains expressing the combination BD alone or fusion, and AD alone or fusion were spotted on SD-Trp-Leu and SD-Trp-Leu-His (in the absence or presence of 3-amino-1,2,4-aminotriazole, 3-AT) plates and incubated until visible colonies were formed. The yeast strain AH109 transformed with vectors pGADT7 + pGBKT7 was used as negative controls.

### Interaction behaviors of PeDL1

Gross *et al.* (2018) reported that CRC could form homodimers and heterodimers with INO. However, our Yeast-two hybrid assays indicated that PeDL1 and PeDL2 do not have the ability to form homodimers. Furthermore, PeDL1 and PeDL2 also could not form heterodimers (Fig. S10a, S10c). To identify transcription factors that interact with PeDL proteins, systematic screening of a yeast-two hybrid transcription factor library composed of approximately 1,350 *Arabidopsis* transcription factors (Mitsuda *et al.*, 2010), was performed with PeDL proteins as preys. The results showed that PeDL1 and PeDL2 were able to interact with several *Arabidopsis* TCP transcription factors (Fig. S9, Tables 4 and 5). We inferred that PeDL1 and PeDL2 might have the ability to form heterodimers with TCP proteins in *Phalaenopsis* orchids. We further examined the protein interactions among PeDLs and *Phalaenopsis* TCP proteins. Three PeCYCs (PeCYC1, PeCYC2, PeCYC3), two PeCINs (PeCIN7, PeCIN8) and four PePCF (PePCF4, PePCF5, PePCF7, PePCF10) were cloned and examined using the yeast two-hybrid system followed by a spotting assay of serial dilutions with cultured yeast cells (Fig. S10). The results showed that PeDL1 could only interact with PeCIN8, whereas PeDL2 could not interact with *Phalaenopsis* TCP proteins tested (Fig. 9, S10). To reconfirm the interaction relationship between PeDL1 and PeCIN8 proteins *in vivo*, the bimolecular fluorescence complementation (BiFC) assay was adopted. The EYFP (enhanced yellow fluorescent protein) signals were detected in PeDL1(EYFP^n^)-PeCIN8(EYFP^c^) as well as PeCIN8(EYFP^n^)-PeDL1(EYFP^c^), revealing that PeDL1 and PeCIN8 could form heterodimers in the nucleus (Fig. 10).

**Fig. 10.**
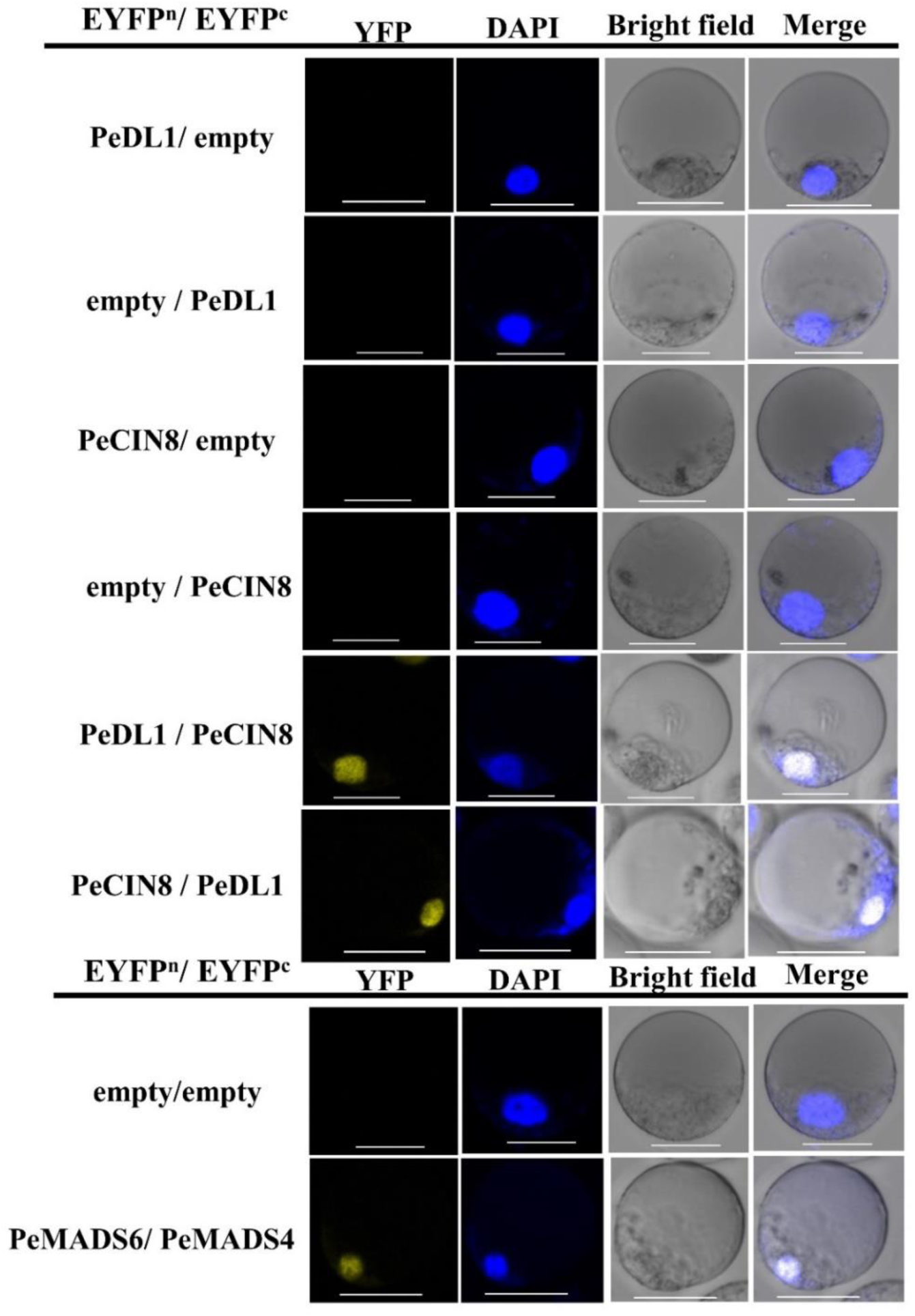
<u>Bi</u>molecular <u>f</u>luorescence <u>c</u>omplementation (BiFC) assay of protein-protein interaction of PeDL1 and PeCIN8 in transiently transfected *P. aphrodite* protoplasts. Combination of non-fused Empty (EYFP^n^) with PeDL1(EYFP^c^) or PeCIN8(EYFP^c^) were unable to reconstitute the fluorescent YFP chromophore. Combination of non-fused Empty (EYFP^c^) with PeDL1(EYFP^n^) or PeCIN8(EYFP^n^) were also unable to reconstitute the fluorescent YFP chromophore. The YFP fluorescence formed through the interaction between PeDL1(EYFP^n^) + PeCIN8(EYFP^c^) or PeDL1(EYFP^c^) + PeCIN8(EYFP^n^) were observed by confocal microscope. Empty (EYFP^n^) + Empty (EYFP^c^) and PeMADS6(EYFP^n^) + PeMADS4 (EYFP^c^) were negative and positive control, respectively. The images were obtained from the YFP fluorescent protein channel, bright field, merged image of the YFP fluorescence, and protoplasts stained with DAPI represented in blue. Scale bars: 20 μm.

## DISCUSSION

Zhang *et al.* (2017) indicated the possibility of an ancient orchid-specific whole-genome duplication (WGD) event shared by all extant orchids, which might be correlated with orchid diversification. Interestingly, it has been reported that DL/CRC represents a single orthologous lineage, without ancient duplications (Lee *et al.*, 2005). Our phylogenetic analysis indicated that orchid DL/CRC could be divided into two subclades. Subclade I includes primitive *Apostasia* DL, *Phalaenopsis* DL1, and *Dendrobium* DL1, whereas subclade II contains *Phalaenopsis* DL2 and *Dendrobium* DL2 (Fig. S1d). The results support that an ancient orchid-specific WGD event generated two *DL/CRC* paralogs in the last common ancestor of orchids, and one of the copies was lost in *Apostasia*, one of two genera that form a sister lineage to the rest of the Orchidaceae. The *Apostasia* might have lost one *DL* gene in subclade II, and duplicated *DL* genes were retained in Epidendroideae orchids. Although *PeDL1* and *PeDL2* have similar expression patterns in reproductive tissues, *PeDL1 per se* presents much higher expression than *PeDL2* in the gynostemium (Fig. 3). Both sequence divergence and differential expression were associated with the functional differentiation of the two *DL/CRC* paralogous genes in *Phalaenopsis*.

The expression patterns of eudicot *DL*/*CRC* genes are consistent with their functions, including termination of the floral meristem, promotion of gynoecium growth, and elaboration of the abaxial carpel wall structures (Yamada *et al.*, 2011). The legumes’ *PsCRC* is not only significantly expressed in carpels through all of the floral buds, but also in the ovary chamber, style–stigma junction, stigmatic tissues, and ovary wall (Fourquin *et al.*, 2014). Some of the core eudicots have recruited *DL*/*CRC*-like genes for nectary development (Lee *et al.*, 2005). In monocot rice, the *DL* is expressed in the floral meristem, whole carpel tissue, and leaf mid-rib (Yamaguchi *et al.*, 2004). Both *PeDL*s’ transcripts detected in the *Phalaenopsis* floral meristem and carpel tissue are consistent with that of ancestral *DL*/*CRC* genes involved in the floral meristem determinacy and carpel specification (Yamada *et al.*, 2011). In addition, expression of both *PeDL*s could be significantly measured in the placenta and the ovule primordia at early stages of ovary development. It is possible that expression of *PeDL*s is acquired in the ovule. In California poppy, *EcCRC* was also independently recruited for additional functions in ovule initiation (Orashakova *et al.*, 2009).

In this study, we provide evidence that overexpression of respective *PeDL1* and *PeDL2* in *Arabidopsis* show loss of inflorescence indeterminacy (Fig. S5). The phenotypes observed agree with *DL*/*CRC* ancestral function in floral meristem determinacy of angiosperms. Interestingly, Wang *et al.* (2011) reported that overexpression of *Cymbidium* orchid *AGAMOUS* (AG) orthologous *CeMADS* genes in *Arabidopsis* also presented similar inflorescence structures. These results were consistent with the fact that *CRC* is directly activated by AG (Gomez-Mena *et al.*, 2005). In *Arabidopsis*, expression of *CRC* driven by its *UBQ10* promoter in *crc-1* mutants could restore the apical fusion of the gynoecium (Gross *et al.*, 2018). To the best of our knowledge, complementation of an *Arabidopsis CRC* gene null mutant by monocot *DL*/*CRC*-like genes has not been achieved so far. Transgenic plants overexpressing respective *PeDL1* and *PeDL2* in *crc-1* demonstrated that both *PeDL*s have conserved functions shared with *CRC* in regulating gynoecium development.

The orchid gynostemium is a unique floral organ, which is fused by three carpels of female and male reproductive organs. The one median carpel is initiated on the abaxial side and two lateral carpel apices protrude on the adaxial side, then the two lateral carpels partly unite with the median carpel, forming the stigmatic cavity (Tsai *et al.*, 2004). The stigmatic cavity provides an elaborate sticky space for accepting pollinium. In our results, transient overexpression of *PeDL1* but not *PeDL2* in *Phalaenopsis* flowers showed altered morphology with protuberances from the bottom of the stigmatic cavity, suggesting functional differentiation of the two paralogous genes. Notably, only one *DL*/*CRC*-like gene was found in the genome of primitive *A. shenzhenica*, which had a gynostemium without stigmatic cavity formations (Kocyan & Endress, 2001). As the ancestral function of *DL*/*CRC* plays a role in the regulation of floral meristem termination and carpel development, one of the duplicated *DL*/*CRC* members in the last common ancestor of orchids might have been co-selected as a regulator of the stigmatic cavity within the Orchidaceae, except for the species in primitive Apostasioideae. *DL*/*CRC*-like genes in monocots are also recruited for additional functions, for example, *DL* has an important function in the differentiation of the Poaceae leaf midrib, different than the ones observed in dicots. Thus, retention of duplicated *DL*/*CRC* genes and functional diversification in the sister lineage to the Apostasioideae could indicate an important role for *DL*/*CRC* genes in orchid evolution.

Protruding tissue growing from the bottom of the stigmatic cavity of transiently overexpressed *PeDL1* indicated that *PeDL1* might regulate the genes involved in cell proliferation. Indeed, comparative transcriptomic analysis revealed that cell cycle related genes are up-regulated in *OE-PeDL1* gynostemium. It is possible that *PeDL1* has the ability to modulate cell proliferation at the adaxial region of the two lateral carpels to contribute to the formation of the stigmatic cavity. In addition, in some *OE-PeDL1* plants, we could observe that the stigmatic cavity was filled with a sticky substance (Fig. S11). Interestingly, several *SWEET* genes with elevated expression in *OE-PeDL1* gynostemium were identified. SWEET has been extensively characterized in plants; these could mediate the sugar transporters (Chen, 2014). For example, *AtSWEET9* is specifically expressed in *Arabidopsis* nectaries and the mutant plant loss of nectar secretion (Lin *et al.*, 2014). *PeDL1* might directly or indirectly promote the expression of *SWEET* genes to create the sticky surface of the stigmatic cavity. Down-regulated *AP2/ERF* and *WRKY* transcription factor genes were also identified from *OE-PeDL1* gynostemium. AP2/ERF and WRKY regulators have major functions for hormone sensitivity and stress responses (Dietz *et al.*, 2010; Banerjee & Roychoudhury, 2015; Phukan *et al.*, 2016; Chandler, 2018; Xie *et al.*, 2019). The results suggest that *PeDLs* not only act as regulators in cell proliferation and sugar transport, but also as modulators of hormone coordination for orchid gynostemium development.

Knowledge of the behavior of YABBY proteins originates from the study of model plants such as *Arabidopsis*, *Antirrhinum*, and rice. In *Arabidopsis*, YAB2, YAB3, YAB5, INO, and CRC can form homodimers (Stahle *et al.*, 2009; Simon *et al.*, 2017; Gross *et al.*, 2018). Heterodimerization was also observed between different YABBY proteins (Stahle *et al.*, 2009; Gross *et al.*, 2018). Interestingly, YABBY proteins could also interact with coactivators as well as corepressors to implement their functions (Liu & Meyerowitz, 1995; Stahle *et al.*, 2009). In *Antirrhinum*, a heterodimer formed by GRAMINIFOLIA (GRAM) and STYLOSA (STY) was described (Navarro *et al.*, 2004). In rice, OsYABBY4 physically interacts with SLENDER RICE 1 (SLR1) to inhibit GA-dependent degradation of SLR1 (Yang *et al.*, 2016). In this study, we observed that both PeDL1 and PeDL2 could not form homodimers and heterodimers with each other (Fig. S10). Interestingly, PeDL1 could interact with type II TCP protein PeCIN8 (Fig. 9 and 10). Alternatively, the expression pattern of *PeDL1* at the early stage of ovule development is parallel with that of *PeCIN8* (Lin *et al.*, 2016). It has been indicated that PeCIN8 plays important roles in orchid ovule development by modulating cell proliferation (Lin *et al.*, 2016). It is possible that the PeDL1-PeCIN8 heterodimer could have functions for the regulation of orchid gynostemium and ovule development. Further study of the downstream targets of PeDL1-PeCIN8 heterodimers may illuminate the molecular basis for unique orchid reproductive development.

## MATERIALS AND METHODS

### Plant Materials

The native species *P. equestris* and *P. aphrodite* subsp. *formosana*, and the *Phalaenopsis* cultivar *P.* Sogo Yukidian were grown in a glasshouse at National Cheng Kung University (NCKU) under natural light (photosynthetic photon flux density, 90 μmol m−2 s−1) and controlled temperature from 23°C to 27°C. The *Arabidopsis thaliana* ecotype *Columbia* (col) and *Landsberg erecta* (Ler) were used in ectopic overexpression and complementation tests, respectively.

### Genome-wide identification of YABBY genes from three sequenced orchid genomes

The conserved YAB domain based on the hidden Markov model (HMM) (PF04690) was obtained from the Pfam protein family database (http://pfam.sanger.ac.uk). To identify the YABBY transcription factor coding genes of orchids, the HMM profile of the YABBY domain was used as a query for an HMMER search (http://hmmer.janelia.org/) of the *P. equestris*, *Dendrobium catenatum*, and *Apostasia shenzenica* genome sequences (E-value=0.00001) (Cai *et al.*, 2015; Zhang *et al.*, 2016; Zhang *et al.*, 2017).

### Sequence alignment and phylogenetic analysis

Multiple sequence alignment of the amino acid sequences of YAB proteins was performed by using ClustalW with default settings. An unrooted Maximum likelihood phylogenetic tree was constructed in MEGA 6 (Tamura *et al.*, 2013) with default parameters. Bootstrap analysis was performed using 1,000 iterations.

### Scanning electron microscopy

The *Arabidopsis* transgenics plant and transient gene overexpression *Phalaenopsis* plants were analyzed by scanning electron microscopy (Hitachi TM3000, Tokyo, Japan), and the samples were frozen in liquid nitrogen. The image data were analyzed using ImageJ software (http://rsb.info.nih.gov/ij/).

### RNA preparation

*Phalaenopsis* orchid samples were collected and stored at −80°C. Total RNA was extracted with the method described by Tsai *et al.* (2004). The five stages of developing flower buds were defined as B1 (0.5-1.0 cm), B2 (1.0-1.5 cm), B3 (1.5-2.0 cm), B4 (2.0-2.5 cm), and B5 (2.5-3.0 cm), based on the description by Pan *et al.* (2014). Definitions of various stages of the developing ovary and ovule from 0 to 64 days after pollination (DAP) were based on the description by Chen *et al.* (2012).

### Quantitative real-time PCR

Total RNA was treated with DNase (NEB, Hertfordshire, UK) to remove remnant DNA. First-strand cDNA was synthesized using the Superscript II kit (Invitrogen, Carlsbad, CA). The quantitative real-time PCR System (ABI 7500, Applied Biosystems) and the SYBR GREEN PCR Master Mix (Applied Biosystems) as described in Lin *et al.* (2016) were used. For each real-time RT-PCR, each sample was analyzed in triplicate. Data were analyzed with the Sequencing Detection System v1.2.3 (Applied Biosystems). *PeActin4* was used as an internal control in Table S6 (Chen *et al.*, 2005).

### RNA *in situ* hybridization

*P. equestris* flower buds, developing ovaries and ovules were fixed in 4% (v/v) paraformaldehyde and 0.5% (v/v) glutaraldehyde for 16-24 hours at 4 °C. Samples were dehydrated with a graded ethanol series (20%, 30%, 50%, 70%, 95%, 100%), and were then sectioned between 6–8 μm with a rotary microtome (MICROM, HM 310, Walldorf). The sense and antisense RNA probes used Digoxigenin-labeled UTP-DIG (Roche Applied Science), and the probe containing a partial C-terminal region was produced by following the manufacturer’s instruction kit (Roche Applied Science). Tissue sections were deparaffinized with xylene, rehydrated through an ethanol series, and pretreated with proteinase K (1 µg mL−1) in 1× PBS at 37°C for 30 min. Pre-hybridization and hybridization followed previous protocols (Tsai *et al.*, 2005).

### Preparation of *Phalaenopsis* petal protoplast

Petal protoplasts were isolated from the fully blooming flowers of *P. aphrodite* subsp. *formosana.* Approximately 5 g of flower buds were sterilized in 70% ethanol for 1 min, followed by three washes in sterilized distilled water. Petals were cut into small pieces and transferred into 8 ml enzyme solution which consisted of 4% cellulose (Onozuka R-10), 2% Macerozyme R-10, 0.6 M sucrose and cell protoplast washing (CPW) salt solution, pH5.8. The digestion was carried out in the dark under gentle shaking at 50 rpm for 4 hours at room temperature. After incubation, the solution was filtered through steel sieves with 100 μm and 50 μm pore sizes to remove the undigested materials. The filtrate was transferred to 15 ml centrifuge tubes and centrifuged at 700 rpm for 5 min. The viable protoplasts floating on the surface were collected. CPW salt solution containing 0.6 M mannitol was used for washing. The purified protoplasts were suspended in 2 ml W5 solution (154 mM NaCl, 125 mM CaCl_2_, 5 mM KCl, 2 mM MES (pH 5.7), 5 mM glucose). The protoplasts were assessed for viability using a haemocytometer and a fluorescence microscope (Leica, DMI 3000B).

### *Arabidopsis* transformation

*PeDL1* and *PeDL2* full-length sequences were cloned into the pBI121 vector, respectively. The plasmids were transformed into *Agrobacterium tumefaciens* (strain GV3101) and the floral dip method was used for *Arabidopsis* transformation (Clough & Bent, 1998). After gene transformation in *Arabidopsis*, we sowed the seeds (T0) on MS media with 50 µg ml^-1^ kanamycin (Sigma-Aldrich). After T0 selection, the kanamycin-resistant seedlings (T1) were transferred to soil and grown as described in Chen *et al.* (2012). In this study, the homozygous lines were used for further analysis.

### Complementation test

To generate *35S-PeDL1 crc*-1 plants, pistils of transgenic *35S-PeDL1* homologous plants were pollinated with pollen grains from *crc-1* homozygous mutants, and all plants in the F1 generation were kanamycin resistant. In the F2 generation, approximately 75% of the plants were kanamycin resistant. For identifying *crc-1* homozygous plants carrying *35S-PeDL1*, genomic DNA was extracted and verified by PCR and sequencing; *35S-PeDL2 crc-1* plants were obtained by adopting the same strategy.

### Transient overexpression of *PeDL*s in *Phalaenopsis* orchid

Full-length coding regions of *PeDL1* and *PeDL2* were respectively cloned into pCymMV expression vectors (Lu *et al.*, 2007). The recombinant vectors were transformed into *Agrobacterium tumefaciens* (strain EHA105). The bacterial cells with pCymMV expression vectors containing *PeDL1* or *PeDL2* were respectively injected into the *Phalaenopsis* inflorescence spike and leaf right above the inflorescence. We generated fifteen independent *PeDL1* and fifteen *PeDL2* transiently overexpressed plants in *P.* Sogo Yukidian. Real-time RT-PCR and next-generation sequencing technology were used to examine the overexpression of *PeDL*s in gynostemium.

### Transcriptome sequencing analysis between gynostemium with *PeDL1* transient overexpression and mock-plant

We extracted the gynostemium total RNA from mock-plants and OE-*PeDL1* plants, then treated it with DNase (NEB, Hertfordshire, UK) to remove genomic DNA. For mock-plant and OE-*PeDL1* plant RNA, three biological replicates were performed. The transcriptome sequencing libraries were prepared with an Illumina HiSeq2000 from Personal Biotechnology Co., Ltd. (http://www.personalbio.cn/). Quality control of the raw sequence data was performed using Fast QC, and hereby filtered to obtain high-quality reads by removing low-quality reads. The high-quality clean reads were mapped to the *P. aphrodite* subsp. *formosana* genome (Chao *et al.*, 2018). We selected those with a fold-change higher than 2 and an adjusted P-value less than 0.05 as the differentially expressed genes (DEGs).

### Yeast two-hybrid assay

For systematically analyzing interaction behavior of PeDLs, *PeDL1* and *PeDL2* were respectively cloned into a pDEST_GBKT7 bait vector, which was obtained from the Arabidopsis Biological Resource Center (ABRC) from the Gateway entry clone. The Y2H Gold yeast strain (Takara Bio Inc.) was employed for the assay. After selecting the yeast harboring bait plasmid, prey plasmids constructed on a pDEST_GAD424 vector (Mitsuda *et al.*, 2010) were additionally transformed and spotted onto positive control medium, which lacks leucine and tryptophan, and test medium, which lacks histidine, leucine, and tryptophan. Yeast growth was observed daily for several days.

For further confirmation of interaction between predicted proteins and PeDLs, Y2H assays were conducted with the MATCHMAKER II system (Clontech, Palo Alto, CA). The target gene full-length sequences were cloned into pGBKT7 bait vectors (GAL4 DNA-binding domain; BD) and pGADT7 prey vectors (GAL4 activation domain; AD), respectively. Two plasmids were co-transformed into AH109 yeast cells and selected on medium lacking leucine and tryptophan (SD-Leu/-Trp, Sigma-Aldrich, St. Louis, MO, USA). The yeast cells were evaluated by serial dilutions and spotting assays. The yeast cells were spotted on SD-Trp-Leu and SD-Trp-Leu-His (in the absence or presence of 3-amino-1,2,4-aminotriazole, 3-AT) plates and incubated until visible colonies were formed. All spotting assay sets were performed as at least three independent experiments.

### Bimolecular fluorescence complementation (BiFC) assay

Gateway-compatible vectors [pSAT5(A)-DEST-cEYFP-N1 and pSAT5-DEST-cEYFP-C1 (Lin *et al.*, 2016)] were used to generate expression vectors by a Gateway cloning strategy. The N-terminus of yellow fluorescent protein (YFP) (YN) was cloned upstream of *PeDL1* and *PeCIN8* in the pE-SPYNE vector, and the C-terminus of yellow fluorescent protein (YC) was fused upstream of *PeDL1* and *PeCIN8* in the pE-SPYCE vector. The signals were visualized by confocal laser microscopy (Carl Zeiss LSM780). Primers used for this study are listed in Table S6.

## SUPPORTING INFORMATION

### Supplemental Figures

**Fig. S1.** Sequence analysis of plant *CRC/DL* genes.

**Fig. S2.** RNA *in situ* hybridization of *PeDL2* longitudinal sections in developing floral buds and cross sections in developing ovules of *P. equestris*.

**Fig. S3.** Comparison of rosette leaves in WT, *35S::PeDL1*and *35S::PeDL2* transgenic plants.

**Fig. S4.** Seed phenotype of WT, *35S::PeDL1* and *35S::PeDL2* transgenic plants.

**Fig. S5.** The inflorescence phenotypes of WT, *35S::PeDL1* and *35S::PeDL2* transgenic plants.

**Fig. S6.** Phenotypes of the gynostemium of mock-treated and OE-*PeDL1* plants.

**Fig. S7.** Phenotypes of the ovary and capsule of mock-treated and OE-*PeDL1* plants.

**Fig. S8.** Volcano plot representation of differential expression genes in the mock-treated and OE-*PeDL1* plants.

**Fig. S9.** Interactome of PeDLs (PeDL1, PeDL2) protein analyzed by efficient Y2H screening using a library composed only of transcription factors in *Arabidopsis*.

**Fig. S10.** Interaction of PeDLs (PeDL1, PeDL2) and with *Phalaenopsis* TCP related-proteins in yeast.

**Fig. S11.** Phenotypes of the gynostemium of mock-treated and OE-*PeDL1* plants in *Phalaenopsis* spp. OX Red Shoe ‘OX1408’.

### Supplemental Tables

**Table. S1.** YABBY-related genes in phylogenetic tree and protein alignment.

**Table. S2.** Up-regulation genes in OE-*PeDL1* gynostemium.

**Table. S3.** Down-regulation genes in OE-*PeDL1* gynostemium.

**Table. S4.** Result of Y2H assay between PeDL1 and *Arabidopsis* TFs.

**Table. S5.** Result of Y2H assay between PeDL2 and *Arabidopsis* TFs.

**Table. S6.** List of primers used in this work.

## ACKNOWLEDGMENT

We thank professor Chiou-Rong Sheue (Department of Life Sciences, National Chung Hsing University) for assisting with the SEM experiment.

## AUTHOR CONTRIBUTIONS

W.-C.T. and Y.-Y.C. designed the research; Y.-Y.C. and Y.-Y.H. performed the research; C.-I.L. and H.-X.Y. conducted transcriptome sequencing and analysis; N.M. conducted Y2H screening and analysis; W.-C.T., Z.-J.L., Y.-Y.C., C.-M.Y., C.-C.C. and S.-B.C. analyzed the data. W.-C.T. and Y.-Y.C. prepared and revised the manuscript.

## REFERENCES

Alvarez J, Smyth DR. 1999. CRABS CLAW and SPATULA, two *Arabidopsis* genes that control carpel development in parallel with AGAMOUS. Development 126(11): 2377–2386.

Banerjee A, Roychoudhury A. 2015. WRKY proteins: signaling and regulation of expression during abiotic stress responses. Scientific World Journal 2015.

Bowman JL, Smyth DRJD. 1999. CRABS CLAW, a gene that regulates carpel and nectary development in *Arabidopsis*, encodes a novel protein with zinc finger and helix-loop-helix domains. Development 126(11): 2387–2396.

Bowman JLJCoipb. 2000. The YABBY gene family and abaxial cell fate. Current Opinion in Plant Biology 3(1): 17–22.

Cai J, Liu X, Vanneste K, Proost S, Tsai WC, Liu KW, Chen LJ, He Y, Xu Q, Bian C, et al. 2015. The genome sequence of the orchid *Phalaenopsis equestris*. Nature Genetics 47(1): 65–72.

Chandler JW. 2018. Class VIIIb APETALA2 ethylene response factors in plant development. Trends in Plant Science 23(2): 151–162.

Chao YT, Chen WC, Chen CY, Ho HY, Yeh CH, Kuo YT, Su CL, Yen SH, Hsueh HY, Yeh JH, et al. 2018. Chromosome-level assembly, genetic and physical mapping of *Phalaenopsis aphrodite* genome provides new insights into species adaptation and resources for orchid breeding. Plant Biotechnology Journal 16(12): 2027–2041.

Chen LQ. 2014. SWEET sugar transporters for phloem transport and pathogen nutrition. New Phytologist 201(4): 1150–1155.

Chen YH, Tsai YJ, Huang JZ, Chen FC. 2005. Transcription analysis of peloric mutants of *Phalaenopsis* orchids derived from tissue culture. Cell Research 15(8): 639–657.

Chen YY, Lee PF, Hsiao YY, Wu WL, Pan ZJ, Lee YI, Liu KW, Chen LJ, Liu ZJ, Tsai WC. 2012. C- and D-class MADS-box genes from *Phalaenopsis equestris* (Orchidaceae) display functions in gynostemium and ovule development. Plant and Cell Physiology 53(6): 1053–1067.

Clough SJ, Bent AF. 1998. Floral dip: a simplified method for *Agrobacterium*-mediated transformation of *Arabidopsis thaliana*. Plant Journal 16(6): 735–743.

Dietz K-J, Vogel MO, Viehhauser A. 2010. AP2/EREBP transcription factors are part of gene regulatory networks and integrate metabolic, hormonal and environmental signals in stress acclimation and retrograde signalling. Protoplasma 245(1-4): 3–14.

Dirks-Mulder A, Ahmed I, Krol L, Menger N, Snier J, van Winzum A, de Wolf A, van’t Wout M, Zeegers J, Butôt R. 2019. Morphological and molecular characterization of orchid fruit development. Frontiers in Plant Science 10: 137.

Finet C, Floyd SK, Conway SJ, Zhong B, Scutt CP, Bowman JL. 2016. Evolution of the YABBY gene family in seed plants. Evolution and Development 18(2): 116–126.

Fourquin C, Primo A, Martínez-Fernández I, Huet-Trujillo E, Ferrándiz C. 2014. The CRC orthologue from *Pisum sativum* shows conserved functions in carpel morphogenesis and vascular development. Annals of Botany 114(7): 1535–1544.

Gomez-Mena C, de Folter S, Costa MM, Angenent GC, Sablowski R. 2005. Transcriptional program controlled by the floral homeotic gene AGAMOUS during early organogenesis. Development 132(3): 429–438.

Gross T, Broholm S, Becker A. 2018. CRABS CLAW acts as a bifunctional transcription factor in flower development. Frontiers in Plant Science 9: 835.

Hsiao YY, Huang TH, Fu CH, Huang SC, Chen YJ, Huang YM, Chen WH, Tsai WC, Chen HH. 2013. Transcriptomic analysis of floral organs from *Phalaenopsis* orchid by using oligonucleotide microarray. Gene 518(1): 91–100.

Hsu H-F, Hsu W-H, Lee Y-I, Mao W-T, Yang J-Y, Li J-Y, Yang C-H. 2015. Model for perianth formation in orchids. Nature Plants 1(5): 15046.

Kocyan A, Endress PK. 2001. Floral structure and development of *Apostasia* and *Neuwiedia* (Apostasioideae) and their relationships to other Orchidaceae. International Journal of Plant Sciences 162(4): 847–867.

Lee JY, Baum SF, Oh SH, Jiang CZ, Chen JC, Bowman JL. 2005. Recruitment of CRABS CLAW to promote nectary development within the eudicot clade. Development 132(22): 5021–5032.

Li H, Liang W, Yin C, Zhu L, Zhang D. 2011. Genetic interaction of OsMADS3, DROOPING LEAF, and OsMADS13 in specifying rice floral organ identities and meristem determinacy. Plant Physiology 156(1): 263–274.

Lin IW, Sosso D, Chen LQ, Gase K, Kim SG, Kessler D, Klinkenberg PM, Gorder MK, Hou BH, Qu XQ, Carter CJ, Baldwin IT, Frommer WB (2014) Nectar secretion requires sucrose phosphate synthases and the sugar transporter *SWEET9*. Nature 508: 546–549

Lin YF, Chen YY, Hsiao YY, Shen CY, Hsu JL, Yeh CM, Mitsuda N, Ohme-Takagi M, Liu ZJ, Tsai WC. 2016. Genome-wide identification and characterization of TCP genes involved in ovule development of *Phalaenopsis equestris*. Journal of Experimental Botany 67(17): 5051–5066.

Liu Z, Meyerowitz EM. 1995. LEUNIG regulates AGAMOUS expression in *Arabidopsis* flowers. Development 121(4): 975–991.

Lu HC, Chen HH, Tsai WC, Chen WH, Su HJ, Chang DC, Yeh HH. 2007. Strategies for functional validation of genes involved in reproductive stages of orchids. Plant Physiology 143(2): 558–569.

Mitsuda N, Ikeda M, Takada S, Takiguchi Y, Kondou Y, Yoshizumi T, Fujita M, Shinozaki K, Matsui M, Ohme-Takagi M. 2010. Efficient Yeast One-/Two-Hybrid Screening Using a Library Composed Only of Transcription Factors in *Arabidopsis thaliana*. Plant and Cell Physiology 51(12): 2145–2151.

Mondragon-Palomino M, Theissen G. 2011. Conserved differential expression of paralogous DEFICIENS- and GLOBOSA-like MADS-box genes in the flowers of Orchidaceae: refining the ‘orchid code’. Plant Journal 66(6): 1008–1019.

Nagasawa N, Miyoshi M, Sano Y, Satoh H, Hirano H, Sakai H, Nagato Y. 2003. SUPERWOMAN1 and DROOPING LEAF genes control floral organ identity in rice. Development 130(4): 705–718.

Navarro C, Efremova N, Golz JF, Rubiera R, Kuckenberg M, Castillo R, Tietz O, Saedler H, Schwarz-Sommer Z. 2004. Molecular and genetic interactions between STYLOSA and GRAMINIFOLIA in the control of *Antirrhinum* vegetative and reproductive development. Development 131(15): 3649–3659.

Orashakova S, Lange M, Lange S, Wege S, Becker A. 2009. The CRABS CLAW ortholog from California poppy (*Eschscholzia californica*, Papaveraceae), EcCRC, is involved in floral meristem termination, gynoecium differentiation and ovule initiation. Plant Journal 58(4): 682–693.

Pan ZJ, Chen YY, Du JS, Chen YY, Chung MC, Tsai WC, Wang CN, Chen HH. 2014. Flower development of *Phalaenopsis* orchid involves functionally divergent SEPALLATA-like genes. New Phytologist 202(3): 1024–1042.

Pan ZJ, Cheng CC, Tsai WC, Chung MC, Chen WH, Hu JM, Chen HH. 2011. The duplicated B-class MADS-box genes display dualistic characters in orchid floral organ identity and growth. Plant and Cell Physiology 52(9): 1515–1531.

Phukan UJ, Jeena GS, Shukla RK. 2016. WRKY transcription factors: molecular regulation and stress responses in plants. Frontiers in Plant Science 7: 760.

Rudall PJ, Bateman RM. 2002. Roles of synorganisation, zygomorphy and heterotopy in floral evolution: the gynostemium and labellum of orchids and other lilioid monocots. Biological Reviews 77(3): 403–441.

Sawa S, Ito T, Shimura Y, Okada K. 1999. FILAMENTOUS FLOWER controls the formation and development of arabidopsis inflorescences and floral meristems. Plant Cell 11(1): 69–86.

Siegfried KR, Eshed Y, Baum SF, Otsuga D, Drews GN, Bowman JL. 1999. Members of the YABBY gene family specify abaxial cell fate in *Arabidopsis*. Development 126(18): 4117–4128.

Simon MK, Skinner DJ, Gallagher TL, Gasser CS. 2017. Integument development in *Arabidopsis* depends on interaction of YABBY protein INNER NO OUTER with coactivators and corepressors. Genetics 207(4): 1489–1500.

Stahle MI, Kuehlich J, Staron L, von Arnim AG, Golz JF. 2009. YABBYs and the transcriptional corepressors LEUNIG and LEUNIG_HOMOLOG maintain leaf polarity and meristem activity in *Arabidopsis*. Plant Cell 21(10): 3105–3118.

Tamura K, Stecher G, Peterson D, Filipski A, Kumar S. 2013. MEGA6: Molecular evolutionary genetics analysis version 6.0. Molecular Biology and Evolution 30(12): 2725–2729.

Tsai W-C, Hsiao Y-Y, Pan Z-J, Kuoh C-S, Chen W-H, Chen H-H. 2008. The role of ethylene in orchid ovule development. Plant Science 175(1-2): 98–105.

Tsai W-C, Kuoh C-S, Chuang M-H, Chen W-H, Chen H-H. 2004. Four DEF-like MADS box genes displayed distinct floral morphogenetic roles in *Phalaenopsis* orchid. Plant and Cell Physiology 45(7): 831–844.

Tsai W-C, Lee P-F, Chen H-I, Hsiao Y-Y, Wei W-J, Pan Z-J, Chuang M-H, Kuoh C-S, Chen W-H, Chen H-H. 2005. PeMADS6, a GLOBOSA/PISTILLATA-like gene in *Phalaenopsis equestris* involved in petaloid formation, and correlated with flower longevity and ovary development. Plant and Cell Physiology 46(7): 1125–1139.

Tsai W-C, Pan Z-J, Su Y-Y, Liu Z-J 2014. New insight into the regulation of floral morphogenesis. International Review of Cell and Molecular Biology 311: Elsevier, 157–182.

Villanueva JM, Broadhvest J, Hauser BA, Meister RJ, Schneitz K, Gasser CS. 1999. INNER NO OUTER regulates abaxial-adaxial patterning in *Arabidopsis* ovules. Genes Development 13(23): 3160–3169.

Wang S-Y, Lee P-F, Lee Y-I, Hsiao Y-Y, Chen Y-Y, Pan Z-J, Liu Z-J, Tsai W-C. 2011. Duplicated C-class MADS-box genes reveal distinct roles in gynostemium development in *Cymbidium ensifolium* (Orchidaceae). Plant and Cell Physiology 52(3): 563–577.

Xie Z, Nolan TM, Jiang H, Yin Y. 2019. AP2/ERF transcription factor regulatory networks in hormone and abiotic stress responses in *Arabidopsis*. Frontiers in Plant Science 10.

Yamada T, Yokota S, Hirayama Y, Imaichi R, Kato M, Gasser CS. 2011. Ancestral expression patterns and evolutionary diversification of YABBY genes in angiosperms. Plant Journal 67(1): 26–36.

Yamaguchi T, Nagasawa N, Kawasaki S, Matsuoka M, Nagato Y, Hirano HY. 2004. The YABBY gene DROOPING LEAF regulates carpel specification and midrib development in *Oryza sativa*. Plant Cell 16(2): 500–509.

Yang C, Ma Y, Li J. 2016. The rice YABBY4 gene regulates plant growth and development through modulating the gibberellin pathway. Journal of Experimental Botany 67(18): 5545–5556.

Zhang GQ, Liu KW, Li Z, Lohaus R, Hsiao YY, Niu SC, Wang JY, Lin YC, Xu Q, Chen LJ, Yoshida K, Fujiwara S, Wang ZW, Zhang YQ, Mitsuda N, Wang M, Liu GH, Pecoraro L, Huang HX, Xiao XJ, Lin M, Wu XY, Wu WL, Chen YY, Chang SB, Sakamoto S, Ohme-Takagi M, Yagi M, Zeng SJ, Shen CY, Yeh CM, Luo YB, Tsai WC, Van de Peer Y, Liu ZJ (2017) The Apostasia genome and the evolution of orchids. Nature 549: 379–383

Zhang GQ, Xu Q, Bian C, Tsai WC, Yeh CM, Liu KW, Yoshida K, Zhang LS, Chang SB, Chen F, et al. 2016. The *Dendrobium catenatum* Lindl. genome sequence provides insights into polysaccharide synthase, floral development and adaptive evolution. Scientific Reports 6: 19029.

